## Supplementary Tables, Figures and Notes for "Mechanistic analysis of enhancer sequences in the Estrogen Receptor transcriptional program"

**Table S1.** List of RF identified important features.

RF important Features

ERα

JUN-1

YBX1

LEF1

NKX3-1

FOXA1

RARα

NR5A2

YBX1:ER

RELA

YBX1:RUNX1

PBX1

PGR

PAX2

NR3C1

RUNX1

**Table S2**. Full table of Literature evidence supporting the roles identified by GEMSTAT models. “% enhancer affected” indicates the percent of enhancers affected by ≥ 5 percentile (ensemble average). “Avg. % change” represents the predicted percentile change in activity after knock down, averaged over enhancers that were affected more than 5 percentile points.

| TF | GEMSTAT inferred role | % enhancer  affected | Avg. % change | Literature-suggested role for TF | Reference PMID |
| --- | --- | --- | --- | --- | --- |
| ERα | Strong activator | 63.8 | -32.5 | Transcriptional activator | - 26884552 |
| FOXA1 | Weak activator | 47.1 | -20.0 | Pioneer factor. Alters DNA accessibility profile and is necessary for Estrogen-induced ERα binding. | - **18358809, 16009131, 21151129** |
| GATA3 | Weak activator | 2.3 | -24.6 | Pioneer factor | - **23172872, 32232341** |
| LEF1 | Weak inhibitor | 2.0 | 13.1 | Represses ERα activity by competitively binding to chromatin/recruiting HDAC1 | 18794125 |
| NKX3-1 | Weak inhibitor | 22.4 | 16.0 | Represses ERα activity by competitively binding to chromatin/recruiting HDAC1 | 18794125 |
| GR | Strong/Weak inhibitor | 0.7 | 18.1 | Unclear; Co-activation of ERα and GR leads to altered ER binding landscape | - **23803465** |
| NR5A2 | Strong activator | 26.7 | -20.2 | Activates ERα-mediated transcription at least partly through co-binding to Estrogen Response Elements. | - **22359603** |
| PAX2 | Strong activator | 7.45 | -24.13 | Upregulated in ER+ breast cancer | - **19005469,22168360** |
| PBX1 | Weak activator | 6.4 | -24.7 | Pioneer factor | - **22125492** |
| PGR | Strong/Weak inhibitor | 6.4 | -9.2 | Unclear/controversial | - **26153859, 28729413, 27885264** |
| RARα | Strong Activator | 3.3 | -21.9 | Unclear/controversial | - **19563758, 20080953, 23868472** |
| RELA (NFκb) | Ambiguous (weak activator/strong inhibitor) | 9.3 | -13.1 | - Transcriptional activation; controversial since ERα is known to repress NFκb transcriptional activity | - **25752574 ,20705611 ,19920189,18703630 ,22963717** |
| RUNX1 | Weak activator | 10.55 | -6.75 | Mediates indirect binding (tethered) of ER to DNA | - **20547749** |
| SP1 | Weak activator | 18.8 | -8.7 | Transcriptional activation in part due to mediation of indirect ERα-DNA binding | - **9328340,11250935,11345900** |
| AP2-γ | Strong Activator | 9.0 | -12.8 | Pioneer factor | - **21572391** |
| YBX1 | Weak/Strong inhibitor | 9.8 | 10.1 | Transcriptional repression through direct interaction with ERα | 29180470 |
| ERα:FOXA1 | Cooperative | 17.8 | -17.7 | Cooperative activity essential for ERα activity | - **18358809, 16009131** - **21151129** |
| ERα:GATA3 | Cooperative | 3.1 | -28.1 | Cooperative activity as a pioneer factor | - **23172872, 32232341** |
| ERα:PBX1 | Cooperative | 0.3 | -10.1 | Pioneer factor | **22125492** |
| ER:PGR | Ambiguous | 0.57 | -11.87 | Controversial | - **26153859,28729413,27885264** |
| ERα:RARα | Cooperative | 1.7 | -17.8 | Controversial | - **19563758,20080953, 23868472** |
| ERα: AP2-γ | Cooperative | 1.3 | -8.9 | Pioneer factor | **21572391** |
| ERα:YBX1 | Competitive | 0.6 | 7.3 | Transcriptional repression through direct interaction with ER | 29180470 |

**Table S3.** List of TFs identified through literature survey and the PubMed ID of their corresponding reference.

| Transcription Factor | PMID |
| --- | --- |
| ERα | 26884552 |
| FOXA1 | 16009131, 26884552, 27791031, 21151129 |
| GATA3 | 23172872, 21878914 |
| NFIB | 29180470 |
| YBX1 | 29180470 |
| AR | 27565181 |
| GR | 29279606 |
| PGR | 28729413, 29435103 |
| PBX1 | 22125492 |
| AP-1 | 19339991, 26906743 |
| AP-2γ | 21572391 |
| SP1 | 15695368 |
| NFκb | 25752574 |
| CEBP | 25752574 |
| RARγ | 20080953, 28977594, 21940749 |
| RXR | 21940749 |
| NKX3-1 | 18794125 |
| LEF-1 | 18794125 |
| NR5A2 (LRH-1) | 24049078 |
| RUNX1 | 20547749 |
| OCT4 (POU5F1) | 27065334 |
| MYC | 21779462 |
| MAX | 21779462 |
| XBP1 | 21297881 |
| PPARG | 23375374 |
| PPARD | 23375374 |
| RARα | 20080953, 19563758 |
| RXRβ | 23375374 |
| NR2F2 | 26894976 |
| NR2C1 | 28087820 |

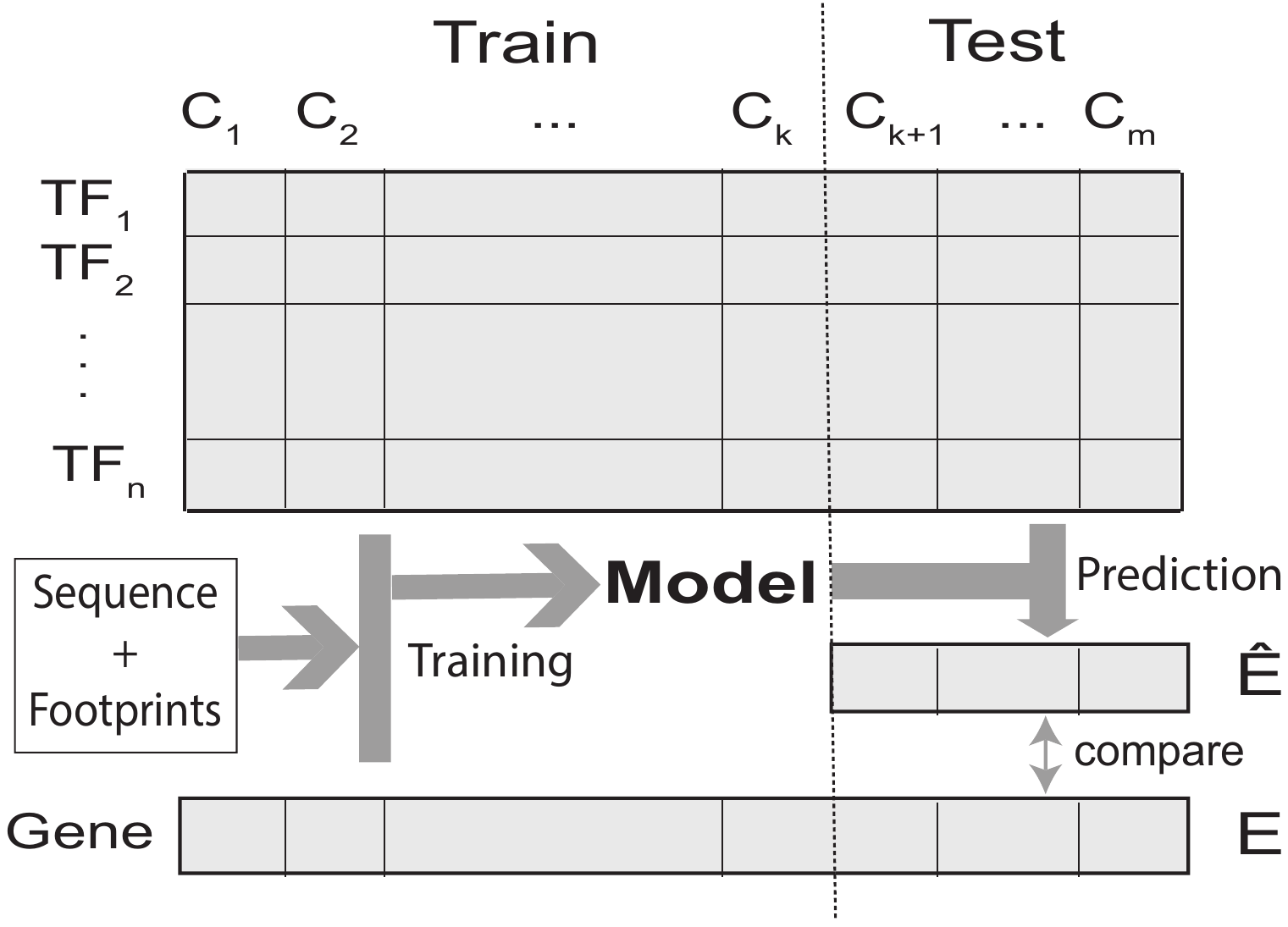

**Figure S1.** Seq2expr Gene expression modeling framework.

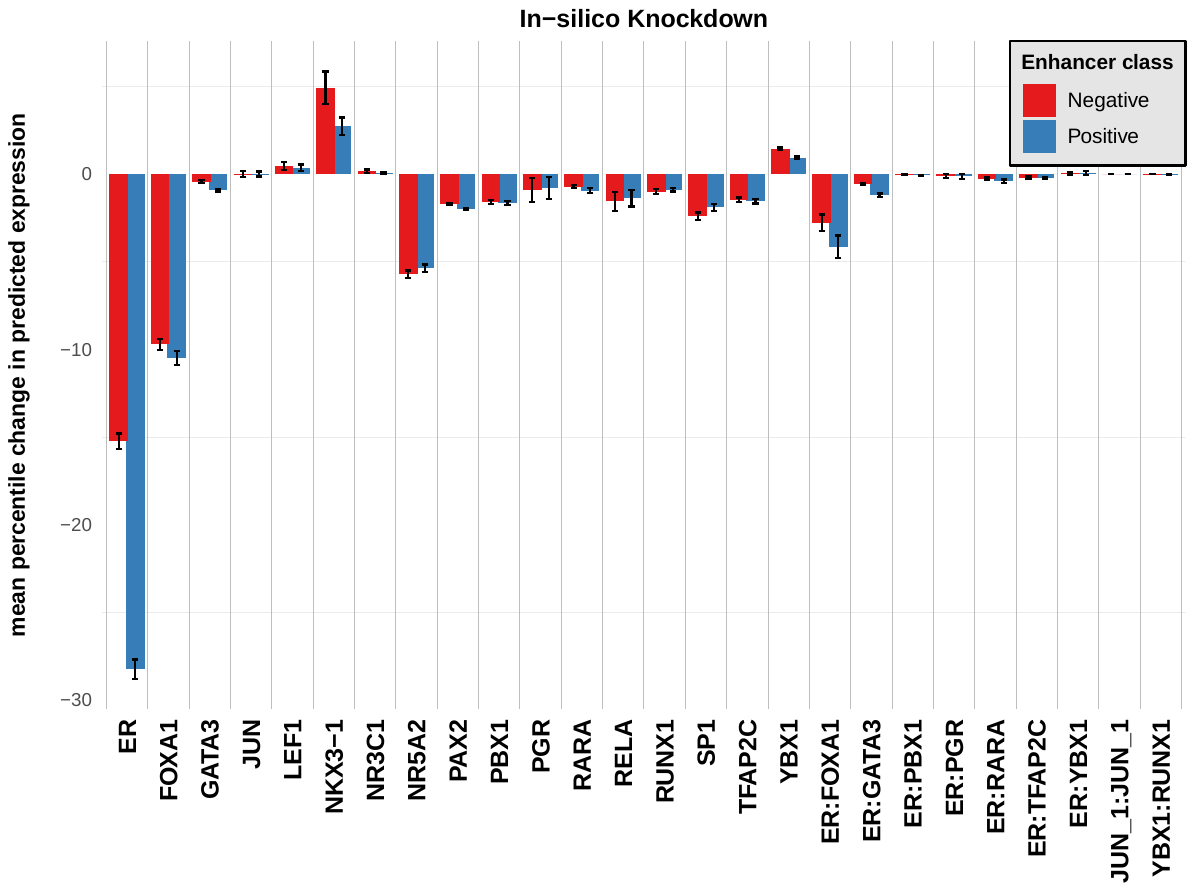

**Figure S2.** In-Silico perturbation of top-ranking ensemble models. Bars represent the perturbation effect of TFs or TF-TF interactions considered by the models, separated by their affecting enhancer class. Positive and Negative classes refer to eRNA expressing and silent enhancers, respectively. Bar height represents the average over 244 top ranking models, and error bars represent the standard deviation. Y-axis indicates the average percentile change in predicted expression due to each trans perturbation.

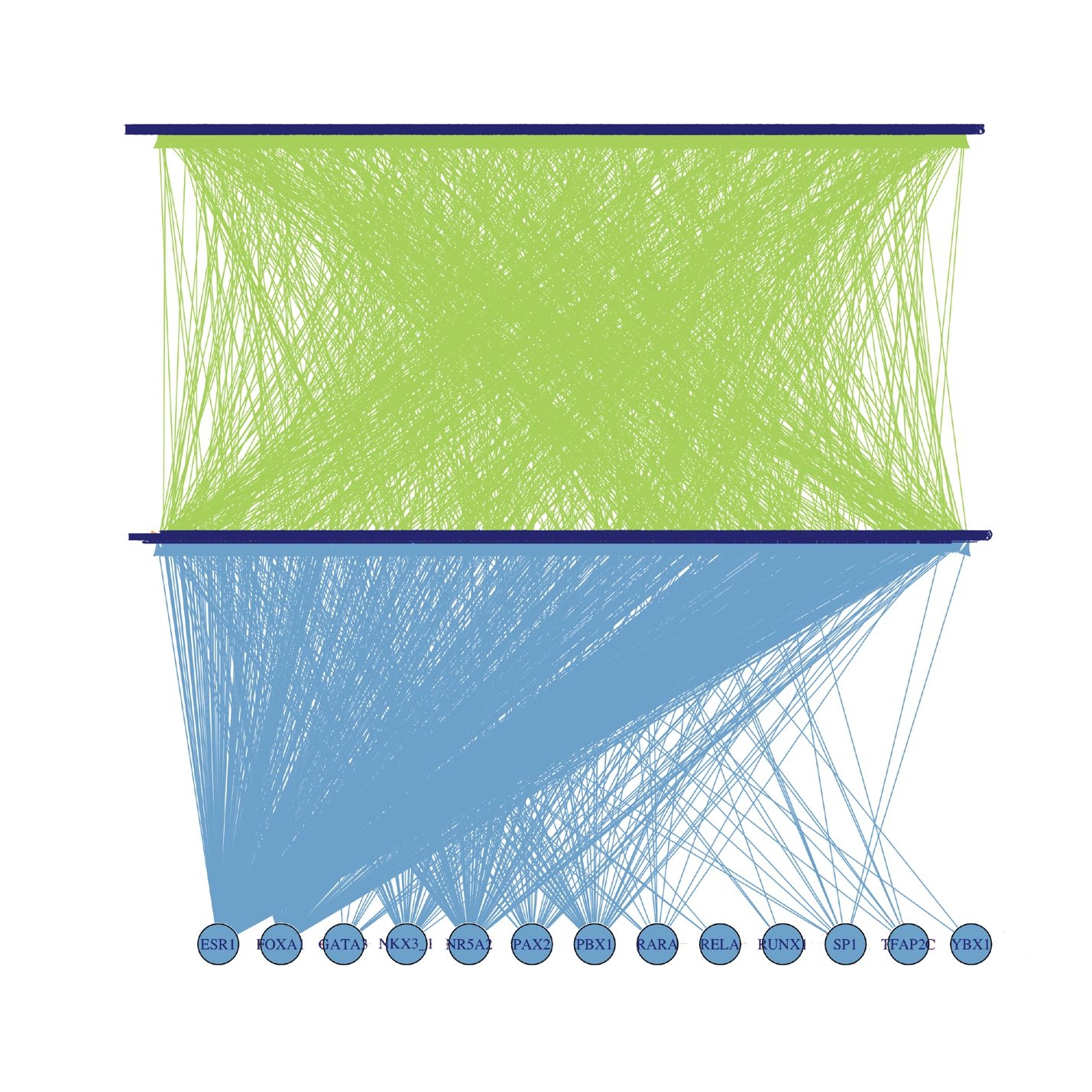

**Figure S3. Overview of TF-enhancer-gene network.** Bottom, middle, and top layers of the network represent TFs, enhancers, and genes, respectively. TF to enhancer edges were derived from in-silico knockdown experiments (Enhancers affected by more than 0.3 percentile levels after a TF knockdown were marked as that TF’s targets.). Enhancer to gene edges represent genomic proximity (+/- 10kb) or interaction (downloaded from 4Dgenome specific to MCF-7 cell line).

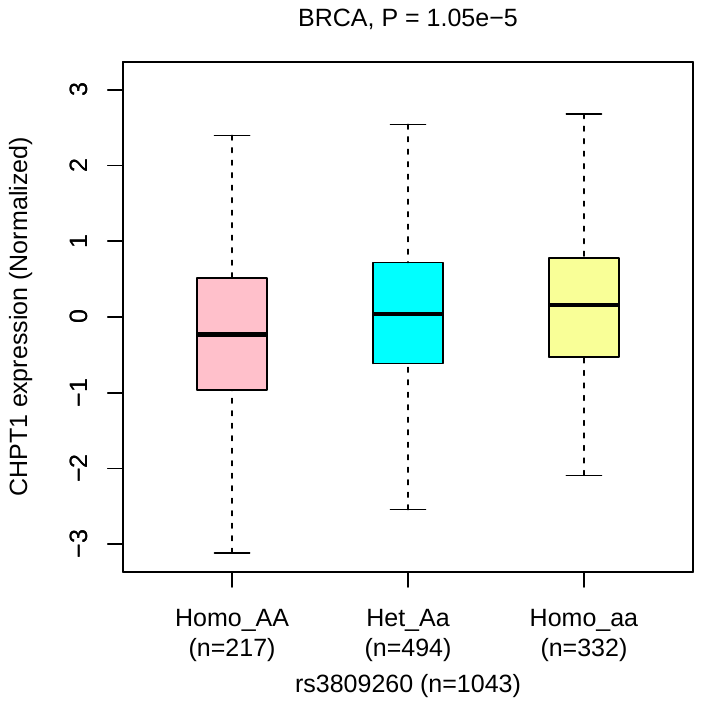

**Figure S4.** Expression profile for gene CHPT1 NOXRED1 stratified by rs3809260 variant in BRCA patients. Figure obtained from PancanQTL.

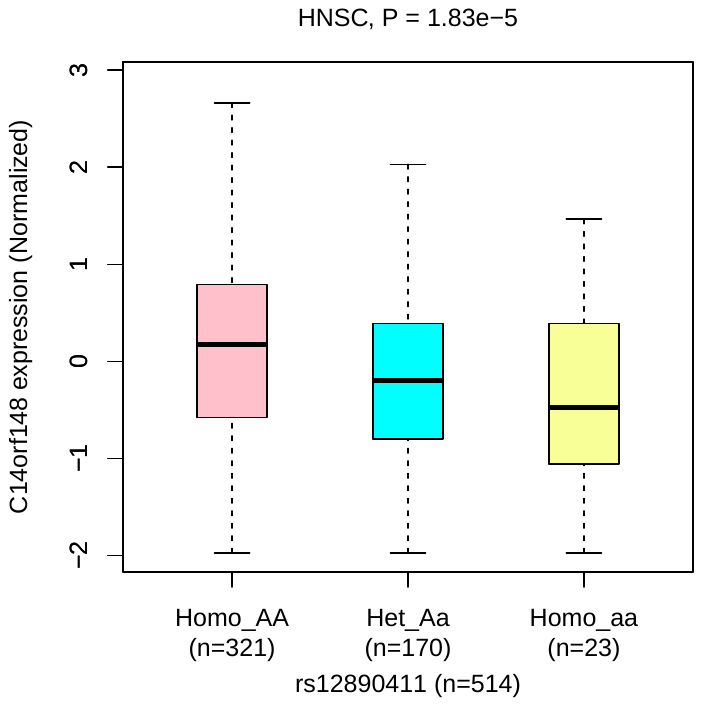

**Figure S5.** Expression profile for gene NOXRED1 stratified by rs12890411 variant in HNSC patients. Figure obtained from PancanQTL.

**Supplementary Note 1**

Genetic variants used in this study were obtained from the following sources:

(1) Breast/Mammary tissue GTEX eQTLs downloaded from https://storage.googleapis.com/gtex_analysis_v8/single_tissue_qtl_data/GTEx_Analysis_v8_eQTL.tar],
(2) Pancan BRCA cis and trans eQTLs downloaded from http://gong_lab.hzau.edu.cn/PancanQTL/static/download/BRCA_tumor.cis_eQTL.xls and http://gong_lab.hzau.edu.cn/PancanQTL/static/download/BRCA_tumor.trans_eQTL.xls
(3) COSMIC breast cancer non-coding variants downloaded from <https://cancer.sanger.ac.uk/cosmic/download> and filtered by cancer type: “breast”
(4) Common SNPs are obtained from dbSNP build 151, subject to a minor allele frequency threshold of greater than 1%. [downloaded from <https://ftp.ncbi.nih.gov/snp/organisms/human_9606_b151_GRCh38p7/VCF/00-common_all.vcf.gz>].

**Supplementary Note 2**

A grid search was performed to identify optimal annotation threshold parameter among three candidates for each TF. In this exercise we trained randomly initialized GEMSTAT ensembles with three different fixed annotation thresholds for each TF, one by one (17 *3 ensembles). The validation performance of GEMSTAT models were used to identify the optimal annotation threshold parameter for each TF. In this exercise we obtained annotation threshold for each of the 17 considered TFs. Additionally, parameter sets from models with best validation performance in this grid search exercise were used as initialization points for the final GEMSTAT ensemble run with optimized annotation thresholds.
